# The Potency of Luliconazole, against Clinical and Environmental *Aspergillus* Nigri Complex

**DOI:** 10.1101/541185

**Authors:** Sahar Hivary, Mahnaz Fatahinia, Simin Taghipour, Marzieh Halvaeezadeh, Ali Zarei Mahmoudabadi

**Affiliations:** Department of Medical Mycology, School of Medicine, Ahvaz Jundishapur University of Medical Sciences, Ahvaz, Iran; Infectious and Tropical Diseases Research Center, Health Research Institute, Ahvaz Jundishapur University of Medical Sciences, Ahvaz, Iran

**Keywords:** Black aspergilli, Luliconazole, Clinical and environmental isolates, Antifungal profile

## Abstract

Black aspergilli are, the most causes of otomycosis and *Aspergillus niger* and *A. tubingensis* are two more frequently isolates. Although, amphotericin B was a gold standard for the treatment of invasive fungal infection for several decades, it replaced by fluconazole and /or voriconazole. Luliconazole, appears to offer the potential for *in vitro* activity against black aspergilli. The aim of the present study was to compare the *in vitro* activity of a novel antifungal agent, luliconazole, with commonly used antifungals against clinical and environmental strains of black aspergilli. Sixty seven strains of black aspergilli were identified using morphological and molecular tests (β-Tubulin gene). Antifungal susceptibility test was applied according to CLSI M38 A2. The results were reported as MIC/MEC range, MIC/MEC_50_, MIC/MEC_90_ and MIC/MEC_GM_. It was found that the lowest MIC range, MIC_50_, MIC_90_, and MIC_GM_ was attributed to luliconazole in clinical strains. *Aspergillus niger* was the common isolate followed by, *A. tubingensis* and 54.1% (clinical) and 30% (environmental) of isolates were resistant to caspofungin. The highest resistant rate was found in amphotericin B for both clinical (86.5%) and environmental (96.7%) strains. Clinical strains of *Aspergillus* were more sensitive to voriconazole (86.7%) than environmental strains (70.3%). On the other hand, 83.8% of clinical and 70% of environmental isolates were resistant to posaconazole, respectively. In conclusion, luliconazole compare to routine antifungals is a potent antifungal for *A. niger* complex *in vitro*. The MIC range, MIC_50_, MIC_90_ and MIC_GM_ of luliconazole against black aspergilli were the lowest among the representative tested antifungals.

## Introduction

Luliconazole (Luzu®), (-)-(E)-[(4R)-4-(2,4-dichlorophe-nyl)-1,3-dithiolan-2-ylidene] (1H-imidazol-1-yl) acetonitrile), is an imidazole antifungal with molecular formula: C_14_H_9_Cl_2_N_3_S_2_ [1]. Luliconazole was basically introduced as anti-dermatophytic antifungal in Japan and India [1, 2]. However, it has demonstrated activity *in vitro* against multiple *Aspergillus* spp. including *Aspergillus fumigatus* [3, 4], *A. terreus* [4, 5], *A. flavus* [4, 6], *A. niger* [4] and *A. tubingensis* [4]. The availability of a novel antifungal, luliconazole, appears to offer the potential for improved therapy for a wide range of invasive fungal infections, including aspergillosis, dermatophytosis, and onychomycosis [2, 7, 8].

While, amphotericin B was a Gold standard in the first-line treatment of invasive fungal infections for several decades [9], it has been replaced by several new antifungals including, voriconazole, posaconazole and caspofungin [10, 11]. Voriconazole was presented as the primary therapy for invasive pulmonary aspergillosis in a clinical trials [12]. Further studies have shown that posaconazole is a useful antifungal for invasive fungal infection including aspergillosis [13]. On the other hand, during 2-3 last decades, caspofungin was developed to improve the prognosis of invasive aspergillosis [14].

The section Nigri (*A. niger*, sensu lato) contains more than 19 accepted species including, *A. niger, A. tubingensis, A. awamory, A. welwitschiae, A. acidus, A. brasiliensis* and others [15-18]. The aspergilli in this section are comprised of several closely related species, and identification based on sequence analyses of β-tubulin gene [4]. *Aspergillus niger* and *A. tubingensis* isolates frequently isolated in clinical infections [16, 19-21]. Black aspergilli cause several types of aspergillosis among predisposed patients [22-25]. Out of them, otomycosis is the most common cutaneous infection caused by black aspergilli [4, 20].

The increasing of fungal opportunistic infections among patients receiving intensive chemotherapy, hematological malignancies and transplant patients during last decades is remarkable [10, 23, 26-28]. Invasive *Aspergillus* infections are one of the life threatening human disease. On the other hand, some species of *Aspergillus* have inherent resistance to some antifungal agents [29]. Moreover, some species have raised minimum inhibitory concentration (MIC) against specific antifungals. As a results, infection prevention consultant and the best choice antifungal are common clinical challenges.

## Objectives

The aim of the present study was to compare the *in vitro* activity of a novel antifungal agent, luliconazole, with amphotericin B, voriconazole, posaconazole and caspofungin against clinical and environmental strains of black aspergilli. Furthermore, the potency of each antifungal against clinical and environmental isolates was compared.

## Materials and Methods

### Fungal isolates

Thirty seven clinical isolates of black aspergilli were previously isolated from otomycosis samples, identified based on morphology characteristics and preserved at Medical Mycology laboratory affiliated to Ahvaz Jundishapur University of Medical Sciences. Environmental strains (30 strains) of black aspergilli were trapped from airborne spores using Sabouraud dextrose agar (SDA) (BioLife, Italia) plates. Primary screening of black aspergilli strains was applied based on macroscopic (Black colony) and microscopic morphology. All strains (clinical and environmental) were subcultured on SDA and re-identified using molecular tests.

### DNA extraction

All strains (clinical and environment isolates) were subcultured on SDA plates and incubated at 29°C for 24 - 48 hours. Mycelia were collected in cryo-tubes containing 300 µL lysis buffer and 0.46 g glass beads and kept at 4°C for 72 hours. The tube contents were homogenized using a SpeedMill PLUS Homogenizer (Analytikjena, Germany) for 6 minutes (3 cycles) and boiled at 100°C for 20 minutes. 300 µL of sodium acetate (3 molar) was added to each tube and stored at -20°C for 10 minutes. Supernatants were removed after a centrifugation at 12000 rpm for 10 minutes. DNA was purified using phenol-chloroform-isoamyl alcohol according to a protocol devised by Makimura et al. [30]. Finally purified DNA was preserved at -20 °C for further tests.

### Molecular identification

β-Tubulin gene was used for the molecular detection of strains using primers pair, βt2a (forward), 5’ GGTAACCAAATCGGTGCTGCTTTC 3’ and βt2b (reverse) 5’ ACCCTCAGTGTAGTGACCCTTGGC 3’ [31]. PCR products subjected for sequence analysis and then sequences were manually verified by MEGA6 software package (https://www.megasoftware.net/) and aligned using the CLUSTALW algorithm. All sequences were compared to reference sequences in the GenBank (NCBI) and CBS database via the nucleotide BLAST ™ algorithm to obtain a definitive identification (similarity values ≥ 99%). Finally, all nucleotide sequences representative were deposited in the GenBank database.

### Antifungal susceptibility assay

Twofold serial dilutions of antifungals including, luliconazole (APIChem Technology, China) (from 0.00012 to 0.25 µg/mL), amphotericin B (Sigma - Aldrich, Germany) (from 0.125 to 16 µg/mL), voriconazole (Sigma - Aldrich, Germany) (from 0.0078 to 4 µg/mL), posaconazole (Sigma - Aldrich, Germany) (from 0.0312 to 4 µg/mL), and caspofungin (Sigma - Aldrich, Germany) (from 0.0078 to 1µg/mL) were prepared in RPMI 1640 (Bio Idea, Iran). Antifungal susceptibility test was performed according to CLSI M38 A2 [32]. A standard suspension (0.5 McFarland) of 48 - 72 hours cultures on SDA was prepared in sterile saline (0.85%) with 0.2% Tween 20 (Merck, Germany). Then, 100 µL of diluted suspension (1:50) and 100 µL of serial dilutions of each antifungal were added to each well of 96-well microplates. Microplates incubated at 35°C for 24 to 72 hours and results were recorded as MIC. Finally, MIC/MEC range, MIC/MEC_50_, MIC/MEC_90_ and MIC/MEC_GM_ were calculated. CLSI or EUCAST have not been defined any clinical or epidemiologic breakpoints/cut offs for amphotericin B, voriconazole, posaconazole, caspofungin and *Aspergillus* species. Strains susceptibility / resistance to each antifungals was evaluated according to commonly utilized breakpoints (Table 1).

**Table 1:** Defined breakpoints of amphotericin B, voriconazole, posaconazole and caspofungin for *Aspergillus niger* sensu lato [33-38].

| Antifungals | MIC/MEC (µg/mL) |  |
| --- | --- | --- |
|  | Sensitive | Resistance |
| Amphotericin B | ≤2 | >2 |
| Posaconazole | ≤0.5 | >0.5 |
| Voriconazole | ≤1 | >1 |
| Caspofungin | ≤0.06 | >0.06 |
| Luliconazole | Undefined | Undefined |

MIC, Minimum inhibitory concentration; MEC, Minimum effective concentration

### Statistical analysis

The Chi-squared test using the Social Science Statistics software (Online) was applied to determine the significant between variables and P value < 0.05 is considered as significance level.

## Results

### Molecular detection of isolates

37 clinical strains of black aspergilli were detected using molecular and sequencing techniques. *Aspergillus niger* (21, 56.8%) was the common strain followed by, *A. tubingensis* (11, 29.8%), *A. luchuensis* (1, 2.7%), and black aspergilli (4, 10.8%) (Table 2). Furthermore, out of 30 environmental black aspergilli isolates, 15 (50%) was identified as *A. niger* followed by, *A. tubingensis* (13, 43.3%), *A. piperis* (1, 3.3%) and black aspergilli (1, 3.3%). However, we could not identified four clinical and one environmental black aspergilli, using molecular technique due to inadequate DNA sample size.

**Table 2:** Clinical and environmental black aspergilli with accession numbers

| Sources | Morphological identification | Molecular identification | Accessions numbers |
| --- | --- | --- | --- |
| Clinical isolates<br>(37 isolates) | *Aspergillus niger*<br>sensu lato | *A. niger*, sensu stricto (21) | LC441155, LC441156, LC441157, LC441158, LC441159, LC441160, LC441161, LC441162, LC441163, LC441164, LC441165, LC441166, LC441167, LC456320, LC456323, LC456326, LC456335, LC456336, LC456337, LC456339, LC456341 |
|  |  | *A. tubingensis* (11) | LC441168, LC441169, LC441170, LC441171, LC456297, LC456298, LC456301, LC456302, LC456303, LC456338, LC456340 |
|  |  | *A. luchuensis* (1) | LC456304 |
|  | Black aspergilli (4) | *** | *** |
| Environmental isolates<br>(30 isolates) | *Aspergillus niger*<br>sensu lato | *A. niger*, sensu stricto (15) | LC456317, LC456318, LC456319, LC456321, LC456322, LC456324, LC456325, LC456327, LC456328, LC456329, LC456330, LC456331, LC456332, LC456333, LC456334 |
|  |  | *A. tubingensis* (13) | LC456299, LC456300, LC456306, LC456307, LC456308, LC456309, LC456310, LC456311, LC456312, LC456313, LC456314, LC456315, LC456316 |
|  |  | *A. piperis* (1) | LC456305 |
|  | Black aspergilli (1) | *** | *** |

### Clinical isolates

It was found that the lowest MIC range (0.00024 - 0.125 µg/mL), MIC_50_ (0.00195 µg/mL), MIC_90_, (0.125 µg/mL) and MIC_GM_ (0.00295 µg/mL) was attributed to luliconazole (Table 3). The minimum effective concentration (MEC) range for all clinical *Aspergillus* species was 0.0078 - 1 μg/ml for caspofungin. In addition, the 50% and 90% MEC (MEC_50_, MEC_90_) values were 0.125 and 0.5 μg/ml for caspofungin. 54.1% of isolates were resistant to caspofungin. The results have shown that the MIC range of amphotericin B for tested isolates was 0.25 - 16 µg/mL. However, MIC_50_, MIC_90_ was similar, 8 µg/mL. The highest resistant rate was found in amphotericin B (86.5%). The MIC ranges for clinical isolates of black aspergilli were 0.0078 - 4 and 0.0625 - 4 µg/mL of voriconazole and posaconazole, respectively. However, the MIC_GM_ for voriconazole (0.77 µg/mL) was lower than posaconazole (1.45 µg/mL). In our study, 29.7% and 83.8% of isolates were resistant to voriconazole and posaconazole, respectively.

**Table 3:** The antifungal susceptibility pattern of 37 clinical strains of black aspergilli

| **Luliconazole** | **N** | **MIC range (µg/mL)** | **MIC<sub>50</sub> (µg/mL)** | **MIC<sub>90</sub> (µg/mL)** | **MIC<sub>GM</sub> (µg/mL)** | **R (%)** |
| --- | --- | --- | --- | --- | --- | --- |
| *Aspergillus niger* | 21 | 0.00024 - 0.125 | 0.00195 | 0.125 | 0.00378 | - |
| *A. tubingensis* | 11 | 0.00024 - 0.125 | 0.00195 | 0.00391 | 0.00251 | - |
| *A. luchuensis* | 1 | 0.00098 | - | - | - | - |
| Black aspergilli | 4 | 0.00049 - 0.00391 | - | - | - | - |
| Total | 37 | 0.00024 - 0.125 | 0.00195 | 0.125 | 0.00295 | - |
| **Amphotericin B** | **N** | **MIC range (µg/mL)** | **MIC<sub>50</sub> (µg/mL)** | **MIC<sub>90</sub> (µg/mL)** | **MIC<sub>GM</sub> (µg/mL)** | **R (%)** |
| *A. niger* | 21 | 0.25 - 8 | 8 | 8 | 4.56 | 17 (81%) |
| *A. tubingensis* | 11 | 4 - 16 | 8 | 8 | 8 | 11 (100%) |
| *A. luchuensis* | 1 | 1 | - | - | - | - |
| Black aspergilli | 4 | 4 - 8 | - | - | - | 4 (100%) |
| Total | 37 | 0.25 - 16 | 8 | 8 | 5 | 32 (86.5%) |
| **Voriconazole** | **N** | **MIC range (µg/mL)** | **MIC<sub>50</sub> (µg/mL)** | **MIC<sub>90</sub> (µg/mL)** | **MIC<sub>GM</sub> (µg/mL)** | **R (%)** |
| *A. niger* | 21 | 0.0625 - 2 | 1 | 2 | 0.99 | 5 (23.8%) |
| *A. tubingensis* | 11 | 0.5 - 4 | 1 | 2 | 1.20 | 4 (36.4%) |
| *A. luchuensis* | 1 | 0.0078 | - | - | - | - |
| Black aspergilli | 4 | 0.5 - 2 | - | - | - | 2 (50%) |
| Total | 37 | 0.0078 - 4 | 1 | 2 | 0.77 | 11 (29.7%) |
| **Posaconazole** | **N** | **MIC range (µg/mL)** | **MIC<sub>50</sub> (µg/mL)** | **MIC<sub>90</sub> (µg/mL)** | **MIC<sub>GM</sub> (µg/mL)** | **R (%)** |
| *A. niger* | 21 | 0.0625 - 4 | 2 | 2 | 1.26 | 17 (81%) |
| *A. tubingensis* | 11 | 0.125 - 4 | 2 | 4 | 2.13 | 10 (90.9%) |
| *A. luchuensis* | 1 | 0.5 | - | - | - | - |
| Black aspergilli | 4 | 0.25 - 4 | - | - | - | 4 (100%) |
| Total | 37 | 0.0625 - 4 | 2 | 4 | 1.45 | 31 (83.8%) |
| **Caspofungin** | **N** | **MEC range (µg/mL)** | **MEC<sub>50</sub> (µg/mL)** | **MEC<sub>90</sub> (µg/mL)** | **MEC<sub>GM</sub> (µg/mL)** | **R (%)** |
| *A. niger* | 21 | 0.0078 - 1 | 0.125 | 0.5 | 0.099 | 11 (52.4%) |
| *A. tubingensis* | 11 | 0.032 - 0.5 | 0.125 | 0.5 | 0.133 | 7 (63.6%) |
| *A. luchuensis* | 1 | 0.032 | - | - | - | - |
| Black aspergilli | 4 | 0.0625 - 0.25 | - | - | - | 2 (50%) |
| Total | 37 | 0.0078 - 1 | 0.125 | 0.5 | 0.107 | 20 (54.1%) |

N, number; MEC, Minimum effective concentration; MIC, Minimum inhibitory concentration; R, Resistant

### Environmental isolates

Table 4 summarizes the *in vitro* susceptibilities of 30 environmental *Aspergillus* Nigri against several antifungals. The same as clinical isolates, the lowest MIC range was 0.00049 - 0.00781 μg/ml for luliconazole. Moreover, the MIC_50_, MIC_90_ and MIC_GM_ were 0.00195, 0.00391 and 0.00195 μg/ml, respectively. The MEC range, MEC_50_, MEC_90_ and MEC_GM_ for caspofungin were 0.0078 - 0.5, 0.0625, 0.25, and 0.0507 μg/ml, respectively. Furthermore, 30% of environmental strains were resistant to caspofungin. As shown, the MIC range for amphotericin B was 2 - 16 μg/ml followed by, MIC_50_, MIC_90_ and MIC_GM_ were 8, 8 and 6.063 μg/ml, respectively. Moreover, 96.7% of strains were resistant to antifungal. Totally, the MIC range voriconazole for environmental isolates of *Aspergillus* was 0.0625 - 2 μg/ml, whereas MIC_90_ 2 μg/ml, MIC_50_ 0.5 and MIC_GM_ 0.4665 μg/ml). Our results indicated that only 4 (13.3%) of strains were resistant to voriconazole. The tested isolates were inhibited at MIC range 0.0625 - 4 μg/ml by posaconazole. Furthermore, the MIC_50_, MIC_90_ and MIC_GM_ were 2, 4 and 1.2599 μg/ml, respectively. In addition, 70% of strains were resistant to posaconazole.

**Table 4:**
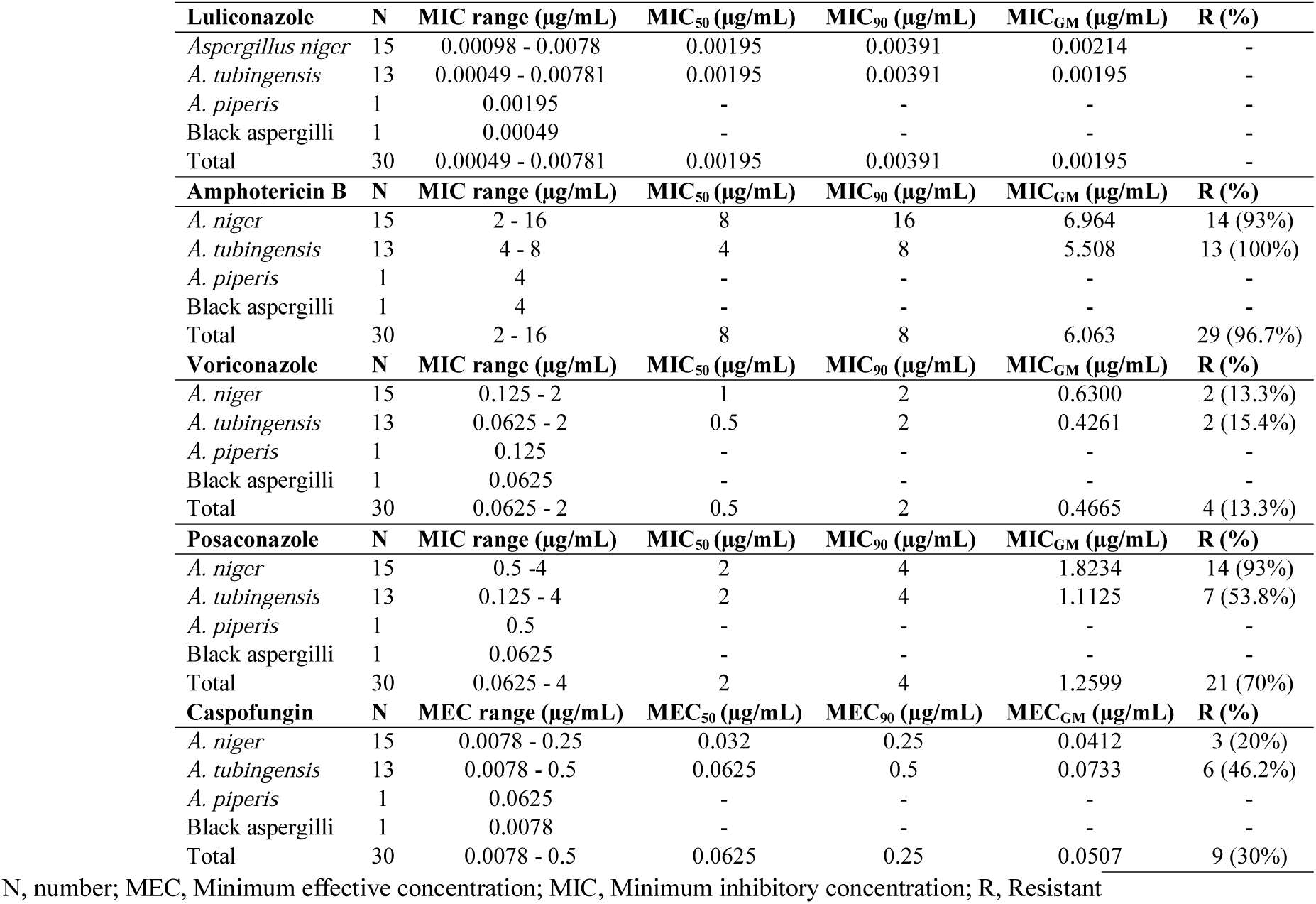
The antifungal susceptibility pattern of 30 environmental strains of black aspergilli

Caspofungin was significantly more effective against environmental than clinical strains (P = 0.048). However, the inhibitory effect of other antifungals (amphotericin B, posaconazole and voriconazole) against both strains (clinical and environmental) was similar (amphotericin B, P=0.147; voriconazole, P=0.109; posaconazole, P=0.178). When we compared the effect antifungals against *A. niger* and *A. tubingensis* among clinical and environmental strains, it is found that caspofungin was more effective on *A. niger* with environmental sources than clinical strains (P=0.0482). Whereas, the effect of other antifungals against both species was not significant.

Our results showed that 32 (86.5%) of clinical strains were resistant to 2, 3 or 4 antifungals, 2 (5.4%) isolates were resistant to one antifungal and 3 (8.1%) were fully susceptible to antifungals (Table 5). Two strains of *A. tubingensis*, one *A. niger* and one black aspergilli were resistant to all antifungals (except luliconazole). On the other hand, in environmental strains, 21 (70%) of strains were resistance to 2 - 4 antifungals and only 30% of strains were resistance to one antifungals (Table 6). Two strains of *A. niger* and one *A. tubingensis* were resistant to all antifungals (except luliconazole).

**Table 5:** Drug resistance against tested antifungals among 37 clinical strains

| Species<br>(Clinical) | Antifungal drugs |  |  |  |  |
| --- | --- | --- | --- | --- | --- |
|  | LUL | POS | VOR | AMP | CAS |
| *Aspergillus niger* | 0.125 | R | S | R | R |
| *A. niger* | 0.125 | S | S | R | R |
| *A. niger* | 0.125 | S | S | R | R |
| *A. niger* | 0.125 | R | S | R | R |
| *A. tubingensis* | 0.125 | R | S | R | R |
| *A. niger* | 0.01561 | R | S | R | R |
| *A. niger* | 0.00781 | R | S | R | R |
| *A. niger* | 0.00781 | R | S | S | S |
| *A. tubingensis* | 0.00391 | R | R | R | R |
| Black aspergilli | 0.00391 | R | S | R | S |
| *A. tubingensis* | 0.00391 | R | R | R | R |
| *A. niger* | 0.00391 | R | S | R | R |
| *A. niger* | 0.00195 | R | R | R | S |
| *A. tubingensis* | 0.00195 | R | R | R | S |
| *A. tubingensis* | 0.00195 | R | R | R | S |
| Black aspergilli | 0.00195 | R | R | R | S |
| *A. tubingensis* | 0.00195 | R | S | R | R |
| *A. niger* | 0.00195 | R | S | R | R |
| *A. tubingensis* | 0.00195 | R | S | R | R |
| *A. niger* | 0.00195 | R | R | R | S |
| *A. tubingensis* | 0.00195 | R | S | R | S |
| *A. niger* | 0.00195 | R | R | R | S |
| Black aspergilli | 0.00195 | R | R | R | R |
| *A. niger* | 0.00195 | R | S | R | S |
| *A. tubingensis* | 0.00195 | S | S | R | R |
| *A. niger* | 0.00098 | R | S | R | S |
| *A. niger* | 0.00098 | R | S | R | R |
| *A. niger* | 0.00098 | R | S | S | S |
| *A. niger* | 0.00098 | R | R | R | R |
| *A. tubingensis* | 0.00098 | R | S | R | S |
| *A. niger* | 0.00098 | R | R | R | S |
| *A. niger* | 0.00098 | R | S | R | R |
| *A. luchuensis* | 0.00098 | S | S | S | S |
| Black aspergilli | 0.00049 | S | S | R | R |
| *A. tubingensis* | 0.00024 | R | S | R | R |
| *A. niger* | 0.00024 | S | S | S | S |
| *A. niger* | 0.00024 | S | S | S | S |

LUL, Luliconazole; POS, Posaconazole; VOR, Voriconazole; AMP, Amphotericin B; CAS, Caspofungin; R, Resistance; S, Susceptible

**Table 6:** Drug resistance against tested antifungals among 30 environmental strains

| Species<br>(Environmental) | Antifungal drugs |  |  |  |  |
| --- | --- | --- | --- | --- | --- |
|  | LUL | POS | VOR | AMP | CAS |
| *Aspergillus niger* | 0.00781 | R | S | R | S |
| *A. tubingensis* | 0.00781 | R | R | R | S |
| *A. niger* | 0.00391 | R | S | R | S |
| *A. niger* | 0.00391 | R | S | R | S |
| *A. niger* | 0.00391 | R | R | R | R |
| *A. tubingensis* | 0.00391 | R | R | R | R |
| *A. niger* | 0.00391 | R | S | R | S |
| *A. niger* | 0.00195 | R | S | R | S |
| *A. tubingensis* | 0.00195 | R | S | R | R |
| *A. niger* | 0.00195 | R | S | R | S |
| *A. tubingensis* | 0.00195 | S | S | R | S |
| *A. niger* | 0.00195 | R | S | R | S |
| *A. niger* | 0.00195 | R | S | S | S |
| *A. tubingensis* | 0.00195 | S | S | R | S |
| *A. niger* | 0.00195 | R | S | R | R |
| *A. tubingensis* | 0.00195 | S | S | R | S |
| *A. tubingensis* | 0.00195 | R | S | R | R |
| *A. tubingensis* | 0.00195 | R | S | R | R |
| *A. tubingensis* | 0.00195 | R | S | R | S |
| *A. tubingensis* | 0.00195 | R | S | R | R |
| *A. niger* | 0.00195 | R | S | R | S |
| *A. tubingensis* | 0.00195 | S | S | R | S |
| *A. piperis* | 0.00195 | S | S | R | S |
| *A. niger* | 0.00098 | S | S | R | S |
| *A. niger* | 0.00098 | R | S | R | S |
| *A. tubingensis* | 0.00098 | S | S | R | S |
| *A. niger* | 0.00098 | R | R | R | R |
| *A. niger* | 0.00098 | R | S | R | S |
| Black aspergilli | 0.00049 | S | S | R | S |
| *A. tubingensis* | 0.00049 | S | S | R | R |

LUL, Luliconazole; POS, Posaconazole; VOR, Voriconazole; AMP, Amphotericin B; CAS, Caspofungin; R, Resistance; S, Susceptible

## Discussion

*Aspergillus* strains isolated from clinical and air borne samples were identified using classical morphological features and molecular methods. Moreover, their susceptibilities to several antifungals including luliconazole, voriconazole, posaconazole, amphotericin B, and caspofungin were assayed. *Aspergillus tubingensis, A. luchuensis* and *A. piperis* were identified as the cryptic species of *A. niger* sensu lato by the sequence analysis of β-tubulin gene. Several reports have shown that *A. niger* is generally as common causative agent of otomycosis and one of the most important agent for invasive aspergillosis [20, 22, 26, 39, 40]. However, new molecular techniques are indicating that this species comprises 19 cryptic species [4, 16, 21].

Some studies have shown a high efficacy of luliconazole against dermatophytes and onychomycosis agents both *in vivo* and *in vitro* [1, 2, 7, 8, 41]. Furthermore, recently a few studies examined the potency of luliconazole against different species of *Candida, A. fumigatus, A. terreus* and *Fusarium* species [5, 6, 42]. However, the potency profile of luliconazole against *A. niger*, complex is unknown. Our results showed that, although the MIC range for strains was extremely low, this range for environmental strains (0.00781-0.00049 μg/ml) was lower than clinical strains (0.125 - 0.00024 μg/ml). As shown in table 5, only five clinical strains (*A. niger* sensu stricto, 4 isolates and *A. tubingensis*, 1 isolate) have a MIC = 0.125 μg/ml. 30/30 (100%) of environmental and 83.8% of clinical strains had the lowest MICs (MICs < 0.00781 μg/ml) against luliconazole. Moreover, the MIC_GM_ for environmental and clinical strains were 0.00195 and 0.00295 μg/ml, respectively. Abastabar et al. [3] and Omran et al. [6] were tested luliconazole against *A. fumigatus* and *A. flavus*, and found that the antifungal has the lowest MICs against *A. fumigatus* (MIC_90_ 0.002 μg/ml) and *A. flavus* (MIC_90_ 0.032 μg/ml), respectively.

There are the limited data in *in vitro* efficacy of antifungals against the black aspergilli both from clinical and environmental sources. While, the clinical and environmental strains had the same MIC ranges for caspofungin, the resistant to antifungal showed the clear differences between clinical and environmental strains (P = 0.048), where the clinical isolates showed higher resistant rate than the environmental strains. In a report by Badali *et al*., only 6.1% of environmental strains of *A. niger* were resistant to caspofungin and all clinical isolates ranged at 0.008–0.063 μg/ml [21].

The *in vitro* activities of posaconazole, voriconazole, and amphotericin B against clinical *Aspergillus* strains have been reported by Arikan *et al*. [10]. They reported that voriconazole was the most active antifungal against *A. niger*. Comparable to our results, voriconazole was more potent than the other tested antifungals (with exception luliconazole) against both clinical and environmental strains. *Aspergillus tubingensis* resistant strains to amphotericin B was very common both in environment and clinical settings, followed by posaconazole, caspofungin, and voriconazole. However, the resistant rate to amphotericin B was lower among environmental than clinical strains. Hashimoto et al., finding suggests that *A. tubingensis* is intrinsically resistant to azole antifungals [15]. Antifungal susceptibility testing of our *A. tubingensis* strains revealed 90.9% and 53.8% of clinical and environmental isolates were resistant to posaconazole.

In conclusion, luliconazole compare to amphotericin B, voriconazole, posaconazole and caspofungin is a potent antifungal for *A. niger* sensu lato *in vitro*. The MIC range, MIC_50_, MIC_90_ and MIC_GM_ of luliconazole against black aspergilli were the lowest among the representative tested antifungals. However, these results suggest luliconazole can be a viable option for the treatment of infections due to black aspergilli and should be further investigated *in vivo*. There is no available systemic formulation of luliconazole and it is strongly suggested that systemic formulation of drug test *in vivo*.

## Acknowledgments

We would like to thank the Infectious and Tropical Diseases Research Center, Health Research Institute, Ahvaz Jundishapur University of Medical Sciences for their support.

## Authors’ Contribution

Study concept and design, Ali Zarei Mahmoudabadi; isolation and preparing clinical and environmental isolates, Marzieh Halvaeezadeh and Sahar Hivary; conducting the experiments, Sahar Hivary; data analysis and interpretation of the results, Ali Zarei Mahmoudabadi, Mahnaz Fatahinia, and Sahar Hivary; drafting of the manuscript, Ali Zarei Mahmoudabadi; Critical editing Mahnaz Fatahinia.

## Funding support

This study was a part of MSc thesis (Sahar Hivary) supported by a grant (No: OG-96148) from the Ahvaz Jundishapur University of Medical Sciences, Ahvaz, Iran.

## Conflict of interest

No conflict of interest declared.

## Ethics statement

This project was approved by the ethical committee of Ahvaz Jundishapur University of Medical Sciences (IR.AJUMS.REC.1396.1066).

